# Modulation of RNA polymerase processivity affects double-strand break repair in the presence of a DNA end-binding protein

**DOI:** 10.1101/2022.02.08.479637

**Authors:** Priya Sivaramakrishnan, Catherine C. Bradley, Irina Artsimovitch, Katelin M. Hagstrom, Laura Deus Ramirez, Alice Xueyin Wen, Matthew Brandon Cooke, Maya Shaulsky, Christophe Herman, Jennifer A. Halliday

## Abstract

Homologous recombination is the predominant double-strand DNA break repair pathway in *Escherichia coli*. A recent report of non-homologous end joining involving the Ku-like Gam protein of bacteriophage Mu (MuGam) has sparked interest in understanding the functions of DNA end-binding proteins in bacteria. MuGam binds to DNA ends, but how it interferes with DNA repair or transcription in live bacteria remains elusive. In *E. coli*, RNA polymerase secondary channel interactors, such as the Gre factors, play a role in coordinating transcription with DNA replication and break repair. Here we show that MuGam inhibits break repair by slowing down DNA resection and impeding recombination in living cells. Loss of GreA restores break repair in the presence of MuGam by allowing for increased DNA resection due to a potential release of MuGam from the DNA. Using MuGam as a DNA break sensor, we found that breaks are generated when translation is inhibited, more so in the presence of GreA, supporting the model where transcription-translation uncoupling increases transcription/replication collisions. Significantly, this work reveals that modulation of RNA polymerase can impact DNA break repair in presence of a Ku-like protein.

## INTRODUCTION

Double-strand DNA breaks (DSBs) are a lethal form of damage that can arise spontaneously as a by-product of cellular growth and metabolism^1^. The two major pathways of DSB repair are homologous recombination (HR) or non-homologous end joining (NHEJ). All organisms that can perform NHEJ share a common feature - the presence of a Ku protein^2^. The first step of NHEJ involves Ku binding to the ends of DSBs and recruiting enzymes to process the terminal ends of the break. This is followed by ligation of the processed ends to reestablish intact DNA. *Escherichia coli* does not have a Ku protein and lacks a canonical NHEJ pathway^2,3^. However, recently it was shown that the Gam protein from phage Mu (MuGam) can promote NHEJ-like ligation of DSB ends in *E. coli* when the bacterial ligase LigA is overexpressed, resulting in repair of broken ends^4^. This study raises questions about the role of MuGam in DSB repair and the utility of Gam as a model for Ku function in bacterial cells.

MuGam shares significant sequence and structural homology with eukaryotic Ku^5^. The main function of MuGam is to protect the phage’s double-strand linear DNA from bacterial nucleases during Mu’s life cycle^6,7^. MuGam binds to double-stranded DNA with blunt or protruding ends, but not to single-stranded DNA or RNA^5^. This feature of MuGam DNA binding has been exploited to visualize the ends of DSBs in *E. coli* and mammalian cells as well as to inhibit repair after CRISPR cutting^8-10^. In *E. coli*, inducible expression of a MuGam GFP fusion (MuGam-GFP) outside of the context of phage infection allows both spontaneous and induced DSBs to be visualized at the single cell, single genomic locus level^8^. MuGam-GFP was able to detect both single- and double-ended DSBs *in vivo*. Expression of MuGam phenocopies a deletion of *recB*^8^, thought to be a consequence of MuGam hindering RecBCD access to double-strand DNA ends (DSEs), where this enzyme complex binds and processes repair intermediates^11^. The RecBCD complex is involved in HR-mediated repair of DSBs and salvage of broken replication forks^12^. RecBCD binds with high affinity to DSEs, resects or degrades DNA and loads the recombination protein RecA onto resected 3’ ssDNA^12^. More recently, using single molecule tracking on assembled DNA curtains, Bhattacharyya et al. showed that MuGam does not block RecBCD binding to DSEs *in vitro*, but rather slows its exonuclease/helicase activity^4^. The extent of this slow down during RecBCD resection on genomic DNA in living bacterial cells is unknown. Furthermore, how the interplay between MuGam and RecBCD binding to DNA ends can affect DSB repair outcomes, particularly by HR, is also unclear.

DNA repair does not occur in isolation on the DNA template. In mammalian cells, mechanisms exist to silence local transcription upon DNA damage through chromatin remodeling^13^. These mechanisms likely evolved to reduce conflicts between RNAP and repair proteins – conflicts that also threaten bacterial cells. In *E. coli*, transcription-coupled repair (TCR) serves to recruit DNA repair proteins to RNA polymerase (RNAP) complexes stalled at helix-distorting and bulky lesions^14^. The stalled RNAPs are in turn removed by one of two helicases – Mfd (classical TCR)^15^ or UvrD in conjunction with the bacterial alarmone ppGpp^16^, allowing repair enzymes to access the lesion. Another class of transcription factors that play important roles in coordinating RNA synthesis with DNA repair/replication conflicts are the RNAP secondary channel interactors - GreA, GreB and DksA. The *E. coli* Gre factors stimulate the intrinsic cleavage activity of RNAP to generate a new 3’ mRNA end, resetting the transcription elongation complex^17,18^. GreA was recently shown to impede DNA break repair by removing backtracked RNAP complexes on DNA, which can potentially be seen as a signal for recombination^19^. DksA functions during both transcription initiation and elongation and plays an important role in the maintenance of genome stability^20-22^ and has been implicated in preventing replication/transcription conflicts triggered during starvation, outcompeting GreA during DNA break repair and R-loop formation^21,23,24^.

In this study, we investigated how the presence of a DNA end-binding protein, MuGam, impacts DNA break repair *in vivo* where other basic cellular processes co-occur on the same DNA template. We were interested in exploring whether MuGam bound to DNA ends would promote RNAP backtracking, and the consequences of this interaction on DSB repair.

## MATERIALS AND METHODS

### Bacterial strains, plasmids and growth conditions

Bacterial strains used in this study are listed in Table S1 and plasmids in Table S2. Bacteria were grown in LB/TB broth or on LB/TB agar plates at 37°C. In the case of the Δ*greA* deletion strain, growth was maintained at 32°C, and cells were transferred to 37°C to match experimental conditions. Antibiotics were added at the following concentrations when necessary – 30 μg/ml kanamycin (kan), 12.5 μg/ml chloramphenicol (cam), 50 μg/ml carbenicillin (CB) and 3.3 μg/ml tetracycline (tet). Doxycycline (dox) was used at the concentrations indicated to induce MuGam expression from the P_N25_-tetO promoter^8^. 0.2% arabinose or 0.2% glucose were respectively added to induce or repress the I-SceI enzyme from the pBAD promoter. Cells were treated with phleomycin (at the indicated concentrations) to induce DSBs. Strain construction was performed using P1 bacteriophage transduction, as described by Miller^25^, and plasmid transformations were performed by electroporation, followed by appropriate selection^26^.

### Survival assays

Semi-quantitative spot assays for viability were carried out by growing strains overnight in LB/TB (with antibiotics when appropriate), followed by diluting cultures and growing them until OD_600_ of approximately 0.4. The cultures were then serially diluted in M9 salts and 5 μl of each dilution was spotted on LB/TB agar plates with and without the treatment to be tested. Quantitative plating efficiencies were determined by serially diluting overnight cultures and spreading appropriate dilutions on LB/TB agar plates (with and without indicated treatment). Colonies were counted after 48 hours. Percent survival was calculated as the number of colonies formed (CFU/ml) with *vs.* without treatment.

### Western blot of MuGam protein levels

Cell lysates were prepared by diluting overnight cultures 1:100 in LB glucose with 0, 100, 200, or 400 ng/mL doxycycline and growing at 37°C. After reaching OD_600_ of 0.1, all treatments were further incubated for 1 hour. For each sample, 1.0 OD_600_ of cells were pelleted, frozen at -80°C to lyse, resuspended in 2X loading buffer with 5% of β-mercaptoethanol, and boiled at 100°C. Samples were run on a 12% SDS-polyacrylamide gel and transferred to a 0.2 μm PVDF membrane using the Trans-Blot Turbo Transfer system (BioRad). The blots were probed with 1:10,000 dilution of rabbit anti-GroEL^27^ or 1:250 mouse anti-Gam^8^ and then probed with goat anti-rabbit (1:10,000 dilution; Invitrogen #A-21244) or goat anti-mouse (1:3000; Invitrogen# A-21236) secondary antibodies and analyzed for immunofluorescence. Blots were imaged (Azure 400, Azure Biosystems) and quantified using Image J.

### Purification of MuGam protein

A bacterial strain containing polyhistidine-tagged MuGam on an expression vector was grown at 32°C till log phase, after which the inducer Isopropyl β-D-1-thiogalactopyranoside (IPTG) was added and incubation continued for 2 hours. The protein was purified as described by Holzinger et al. with the modifications described as follows^28^. The sonication step was omitted, and cobalt was used to bind the His-tagged protein in a sepharose column. The column was washed with buffer multiple times with serially lowered urea concentrations, and finally the protein was eluted in 2M urea. Dialysis was performed to replace the urea-containing buffer with a solution containing potassium chloride, glycerol and dithiothreitol. MuGam is a 21 kDa protein. Both monomeric and dimeric forms of the protein were visible on a stained protein gel. Protein concentration was estimated by the standard Bradford assay.

### *In vitro* assay for RNAP-Gam interaction

Purified biotin-tagged *E. coli* RNAP was bound to magnetic streptavidin coated beads and allowed to initiate transcription on a blunt-ended DNA template with ^32^P-labeled 5’ end with a subset of NTP substrates, resulting in a halted elongation complex. MuGam and or GreB variant proteins (or storage buffer) were incubated with the halted complex, followed by addition of all NTPs to allow RNAP complex to complete transcription of the template DNA at 37°C. The supernatant, containing the released radiolabeled DNA, was collected at various times after resuming transcription and analyzed by scintillation counting.

### XO-seq

XO-seq was performed as previously described^19^. TB media containing 100 ng/mL dox was inoculated 1:100 with overnight culture and grown at 37°C to OD_600_ ∼0.1. After reaching logarithmic growth, the 0-hour samples were collected and 0.2% arabinose was added to induce I-SceI and cutting. Samples were collected 2 hours after arabinose induction and DNA isolation was performed using the Promega Wizard Genomic DNA Purification kit. The Nextera XT protocol and sample preparation kit was used to prepare libraries for sequencing and alignment on Illumina MiSeq. The resulting BAM files were loaded in Seqmonk software for further analysis. Reads were obtained by generating a set of running windows of 1 kb and a step size of the same length. Reads were corrected for the total read count to per million reads, counting duplicated reads once. In R, reads were log2-transformed, normalized to paired 0 min controls, and graphed, as per the previously published XO-seq method^19^.

### MuGam-GFP foci analysis of dox treated cells

WT *E. coli* containing a dox-inducible MuGam-GFP construct were grown overnight at 32°C in LB, diluted 1:100 in TB glucose and incubated with shaking for 1 hr at 37°C. Cultures were then treated with 100 ng/ml anhydrotetracycline (ATC) plus either 0, 100, 200 or 400 ng/ml dox for 24 hours. ATC has no antibacterial activity and induces the P_n25-_tetO-gam reporter without toxicity. After treatment, cells were washed twice in 1 X MinA salts, fixed in 2% paraformaldehyde for 30 minutes on ice, then washed twice before imaging.

To follow cell growth and MuGam-GFP foci formation *in vivo* over time, 1% TB glucose agar pads were prepared containing dox (400 ng/ml) using Gene Frames (Thermo Scientific #AB-0577). The solidified agar pad was cut in two sections. Following overnight MuGam-GFP induction, 2 µl of WT and Δ*greA* were placed on the separated agar sections and growth of microcolonies at 35°C was followed by time-lapse microscopy. Imaging was performed using a Zeiss HAL100 inverted fluorescence microscope. Fields were acquired at 100X magnification with an sCMOS camera (Photometrics). Bright field and fluorescence images were acquired using Zen.1 (Zeiss) software. Foci analysis, cell counting, and cell length measurements were measured using FIJI.

### Mutation rate analysis

For fluctuation assays, bacteria were grown in TB media and all diluted in 1X MinA salts with 1 mM MgSO_4_. Cultures were grown at 37 °C to saturation (16 hr) and used to seed cultures without or with dox at 200 ng/mL at a cell density of 60 cells/ml. After 40 hours, 500 μl of culture was pelleted and resuspended in MinA salts with 1mM MgSO4 before plating on LB plates containing 50 μg/ml carbenicillin or 50 μg/ml rifampicin. Cultures were also serial diluted and tittered on LB plates. Antibiotic resistant colonies were counted after 48 hours of growth and normalized to total cell counts.

### Statistics

For survival tests, the Kruskal–Wallis test followed by pairwise Wilcox test with multiple testing correction was performed. For XO-seq analysis, two-tailed two sample *t*-tests was used. For quantification of MuGam-GFP foci, a One-way ANNOVA followed by Tukey Posthoc was used. For fluctuation analysis, Welch’s one-sided t-test was performed.

## DATA AVAILABILITY

Source data for figures will be provided on publication. Sequencing data will be available through the Sequence Read Archive under accession number XXXX. All other data are available from the corresponding author upon reasonable request.

## RESULTS

### MuGam induction is lethal in the presence of chronic DNA breaks

The ability of MuGam to phenocopy DNA end-repair deficient cells^8^, in conjunction with its DNA end-binding properties^5^, suggests that MuGam confers cell toxicity by physically blocking access of repair proteins to the DNA. Therefore, the effect of MuGam on DSB repair *in vivo* can be assessed by measuring viability in the presence of DSB-inducing agents. To test this, we treated *E. coli* expressing MuGam from a doxycycline (dox)-inducible promoter with phleomycin (PHL), a drug known to induce DNA breaks and measured survival in LB media (Fig. 1a, 1b and see methods). Similar to what was previously observed^8^, induction of MuGam for ∼48 hours led to a decrease in viability in wild-type (WT) *E. coli* grown in LB (Fig. 1b). In the absence of MuGam, PHL treatment leads to a 5-fold loss of cell viability in completely WT *E. coli* (Fig. 1b). However, when cells containing the inducible MuGam construct are exposed to both PHL and dox, we observed a synergistic 4-log decrease in viability (compared to the same strain with dox alone), suggesting that MuGam is severely toxic in the presence of DNA breaks, likely due to an effective block of DNA repair function (Fig. 1b).

**Figure 1.**
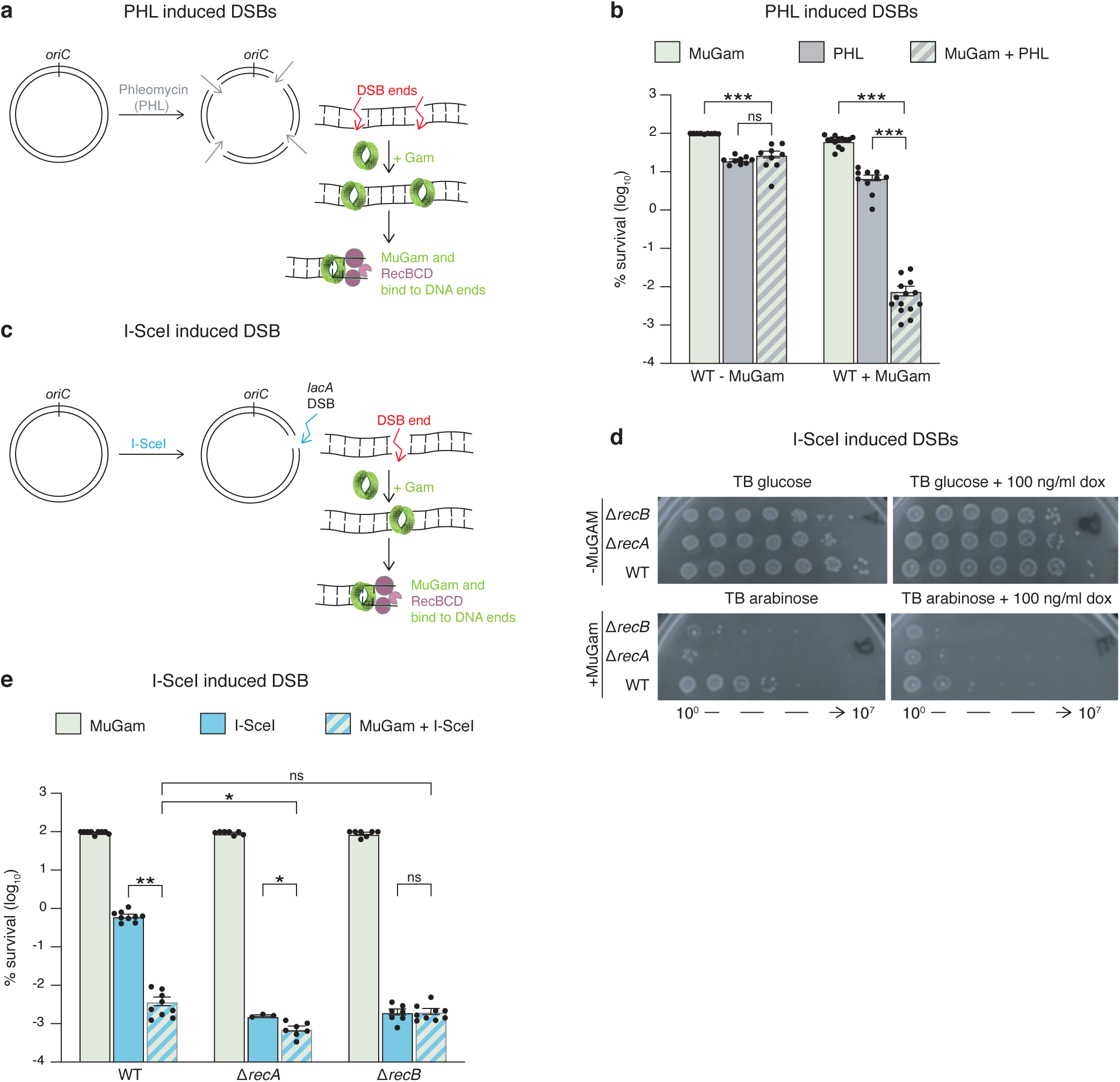
*E. coli* are sensitive to MuGam expression when DSBs are induced *in vivo*. **a** Schematic showing phleomycin (PHL)-induced DSBs bound by MuGam followed by the loading of the RecBCD complex in *E. coli*. **b** Quantitative survival of the wild-type (WT) strain treated with 1 µg/ml PHL with or without 100 ng/ml doxycycline (dox) to induce MuGam expression, mean ± S.E.M. \*\*\**P* ≤ 0.001; \*\**P* ≤ 0.01; \**P* ≤ 0.05; ns, *P* ≥ 0.05 (Kruskal–Wallis test followed by pairwise Wilcox test with multiple testing correction). **c** Expression of the I-SceI enzyme creates a single DSB at *lacA*. **d** Representative viability of the indicated strains without (TB glucose + 100 ng/ml dox) and with I-SceI induced DSBs (TB arabinose + 100 ng/ml dox). **e** Quantitative survival of the indicated mutants with expression of MuGam (100 ng/ml dox), I-SceI induction (0.2% arabinose) or MuGam and I-SceI (100 ng/ml dox + 0.2% arabinose), mean ± S.E.M. \*\*\**P* ≤ 0.001; \*\**P* ≤ 0.01; \**P* ≤ 0.05; ns, *P* ≥ 0.05 (Kruskal–Wallis test followed by pairwise Wilcox test with multiple testing correction).

PHL is a radiomemetic drug that creates DSBs at random positions in the genome. To determine the effects of MuGam when DSBs are generated at a single locus in the *E. coli* genome, we measured survival after DSB induction using an engineered recognition site for the I-SceI endonuclease at the *lacA* locus (Fig. 1c, 1d). I-SceI cutting was performed in TB media, where fewer chromosomal copies are present and repeated cutting at the same site across sister chromosomes impedes homology directed repair. I-SceI DSB induction at the *lacA* locus results in 100-fold loss of survival compared to WT strains lacking the enzyme^19^. When I-SceI cutting is combined with MuGam expression, we observe a ∼5-log decrease in cell viability compared with MuGam expression alone (Fig. 1d,1e).

To confirm that expression of MuGam phenocopies loss of DNA break repair by the canonical HR pathway, Δ*recB* and Δ*recA* null mutations (complete loss of function) were introduced and tested in the I-SceI DSB model. In the presence of MuGam alone, viability of Δ*recA* or Δ*recB* bacteria were similar to WT (Fig. 1d,1e). Both Δ*recA* and Δ*recB* mutants were more sensitive to I-SceI induction alone, but only the Δ*recA* mutant showed significantly lower survival compared with WT in the presence of both MuGam and I-SceI (Fig. 1e). These results confirm that MuGam inhibits DSB-dependent DNA end-repair through HR.

Overall, the two break-induction methods provide complementary approaches to explore the complex processes that occur at DNA ends in presence of MuGam.

### MuGam toxicity is influenced by GreA, a RNAP secondary channel interactor

We previously showed that removal of GreA, which rescues backtracked RNAP complexes (Fig. 2a), increases DSB repair activity upon PHL treatment or I-SceI break induction^19^. To determine if GreA would also influence DSB repair when MuGam is expressed, we first measured cell viability in the presence of both MuGam and PHL in Δ*greA* mutants. The loss of viability observed in WT cells expressing MuGam and treated with PHL is almost completely rescued by the removal of GreA (Fig. 2b). This rescue is dependent on the canonical DSB resection pathway involving RecB (Fig. 2c). As GreA is an auxiliary transcription factor, one possible explanation could be that the absence of GreA reduces MuGam expression levels, thus ameliorating its toxic effects. To rule this out, we performed a Western blot using a MuGam antibody and found no difference in protein levels between WT and the *ΔgreA* cells (Fig. 2d).

**Figure 2.**
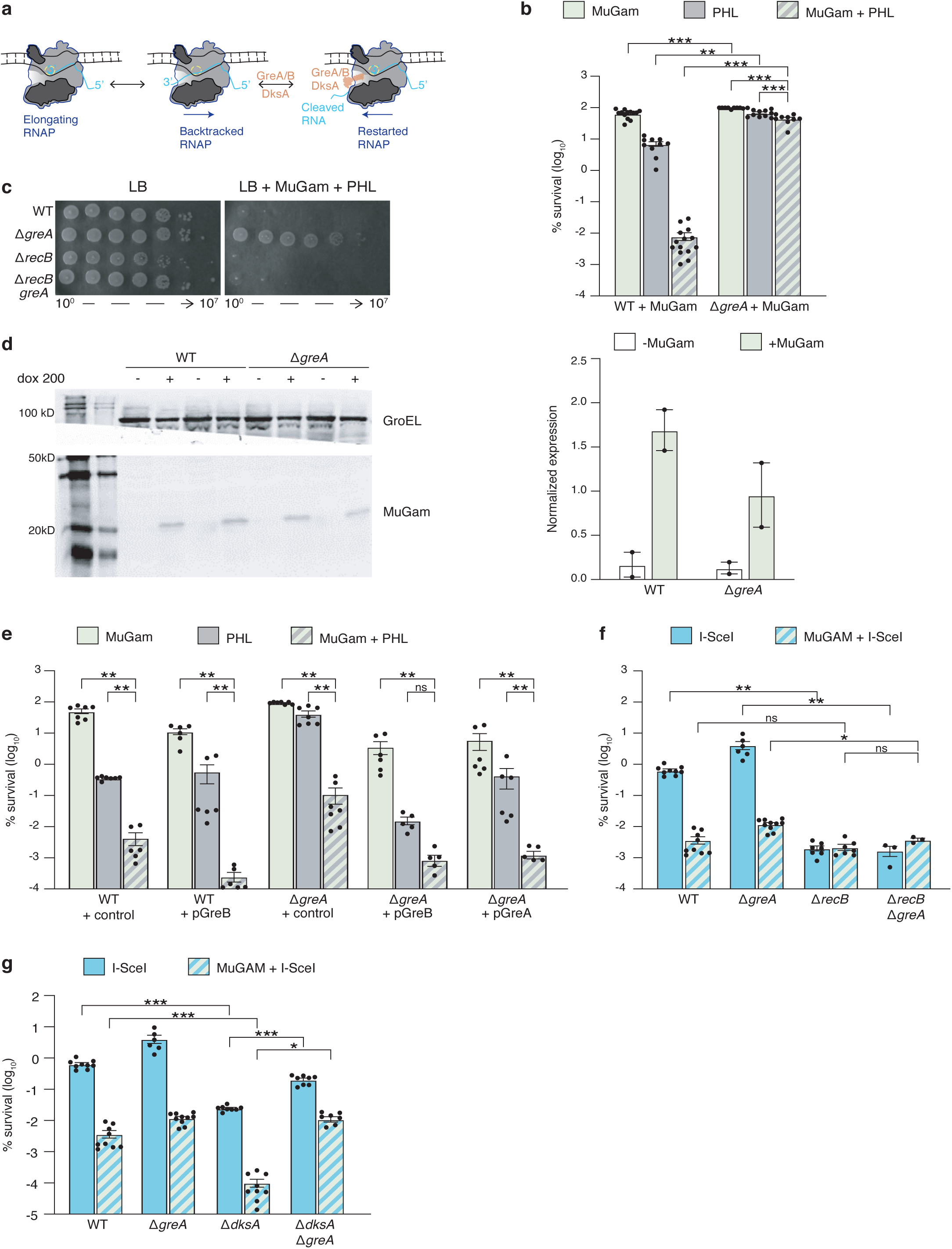
RNAP-secondary channel interactors influence MuGam survival after DSB induction. **a** Schematic showing that GreA/GreB reverse RNAP backtracking by facilitating cleavage of RNA extruded into the secondary channel. DksA also binds to the RNAP secondary channel and can rescue backtracked RNAP without transcript cleavage. **b** Quantitative survival of the indicated mutants treated as in Fig. 1b, mean ±S.E.M.***P<0.001 \*\**P* ≤ 0.01; \**P* ≤ 0.05; ns, *P* ≥ 0.05 (Kruskal–Wallis test followed by pairwise Wilcox test with multiple testing correction). **c** Representative viability of the indicated strains to PHL-induced DSBs with MuGam expression (100 ng/ml dox) or without. **d** Western blot for MuGam in the induced (+dox) and un-induced (-dox) conditions using GroEL for normalization of protein levels and quantification of protein levels, (2 biological replicates) **e** Quantitative survival of 1 µg/ml PHL-induced DSBs in the presence of MuGam (100 ng/ml dox) after expression of control plasmid or high copy plasmid expressing either GreA or GreB with 0.1 mM IPTG in the indicated mutants, mean ±S.E.M. \*\**P* ≤ 0.01; ns, *P* ≥ 0.05 (Kruskal–Wallis test followed by pairwise Wilcox test with multiple testing correction). **f,g** Quantitative survival of the indicated mutants as in Fig. 1e, mean ±S.E.M. \*\*\**P* ≤ 0.001; \*\**P* ≤ 0.01; \**P* ≤ 0.05 ; ns, *P* ≥ 0.05 (Kruskal–Wallis test followed by pairwise Wilcox test with multiple testing correction).

To determine if the loss of viability upon MuGam induction could be affected by other RNAP secondary channel interactors, we tested the effects of GreB, a GreA homolog^18,29^. *E. coli* GreA, an abundant primary cleavage factor, can be functionally replaced by GreB overexpressed from a plasmid^30^. We found that expression of GreB abrogated the resistance of the Δ*greA* mutant to PHL with and without MuGam expression, and further increased MuGam-associated PHL toxicity observed in the WT strain (Fig. 2e). This suggests that both Gre factors can promote MuGam-associated toxicity, consistent with their shared function in stimulating 3’ transcript cleavage to rescue RNAP backtracking^29,31^.

To test whether the GreA effect can be extended to a single DSB, we used the I-SceI system. A milder but significant 4-fold rescue was observed upon MuGam induction in Δ*greA* bacteria exposed to continuous I-SceI cutting compared to a ∼6.5-fold rescue without MuGam induction (Fig. 2f, 2g). As with PHL, the increased survival of the Δ*greA* strain to MuGam and I-SceI expression was dependent on *recB,* suggesting involvement in the canonical RecBCD-dependent recombinational repair pathway (Fg. 2f).

Bacterial Gre factors bind in the RNAP secondary channel and stimulate nascent RNA cleavage in backtracked complexes, thus promoting transcription elongation^17,18,32^ (Fig. 2a). DksA is another RNAP secondary channel interactor that acts as a transcription initiation factor and an elongation factor in living bacterial cells^20,23^. DksA can compete with the Gre factors for RNAP binding^33^ and we have previously described a mechanism where competition between DksA and GreA for RNAP binding influences transcription-DNA break repair conflicts^19^. However, DksA likely has additional roles in DNA repair, which might depend on the type of lesion and damaging agent^22^. While the Δ*dksA* strain was much more sensitive to MuGam expression with I-SceI DSB induction than WT cells, deletion of *greA* restored Δ*dksA* viability (Fig. 2g), consistent with the competition between GreA and DksA for the secondary channel.

Altogether, these results suggest that modulation of RNAP by secondary channel interactors affects the activity of MuGam at DNA breaks.

### DNA resection at DSBs is altered by MuGAM expression

We next looked at the pattern of DNA resection during break repair in the presence of MuGam to examine repair kinetics in live bacteria. DSB repair in *E. coli* proceeds through dynamic intermediates of RecBCD-driven DNA resection and RecA-mediated recombinational repair^12,34^. Capturing these intermediates *in vivo* has historically been a technical hurdle for the DNA repair field - a problem which XO-seq was developed to address^19^. In XO-seq, following induction of a DSB, resection at the breakpoint locally depletes sequencing coverage, which is then reconstituted by recombination-mediated DNA synthesis. By graphing read counts across cutsite-proximal loci, it is possible to visualize the average of resection and recombination activity. In WT cells, read loss after I-SceI cutting at the *lacA* locus is asymmetrical around the break site, extending less than 100 Kb in the origin-proximal direction, but more than 200 Kb on the other side of the break (Fig. 3a). This asymmetry can be attributed to the different distribution of active Chi sites upstream and downstream of the cut-site^12,19^.

**Figure 3.**
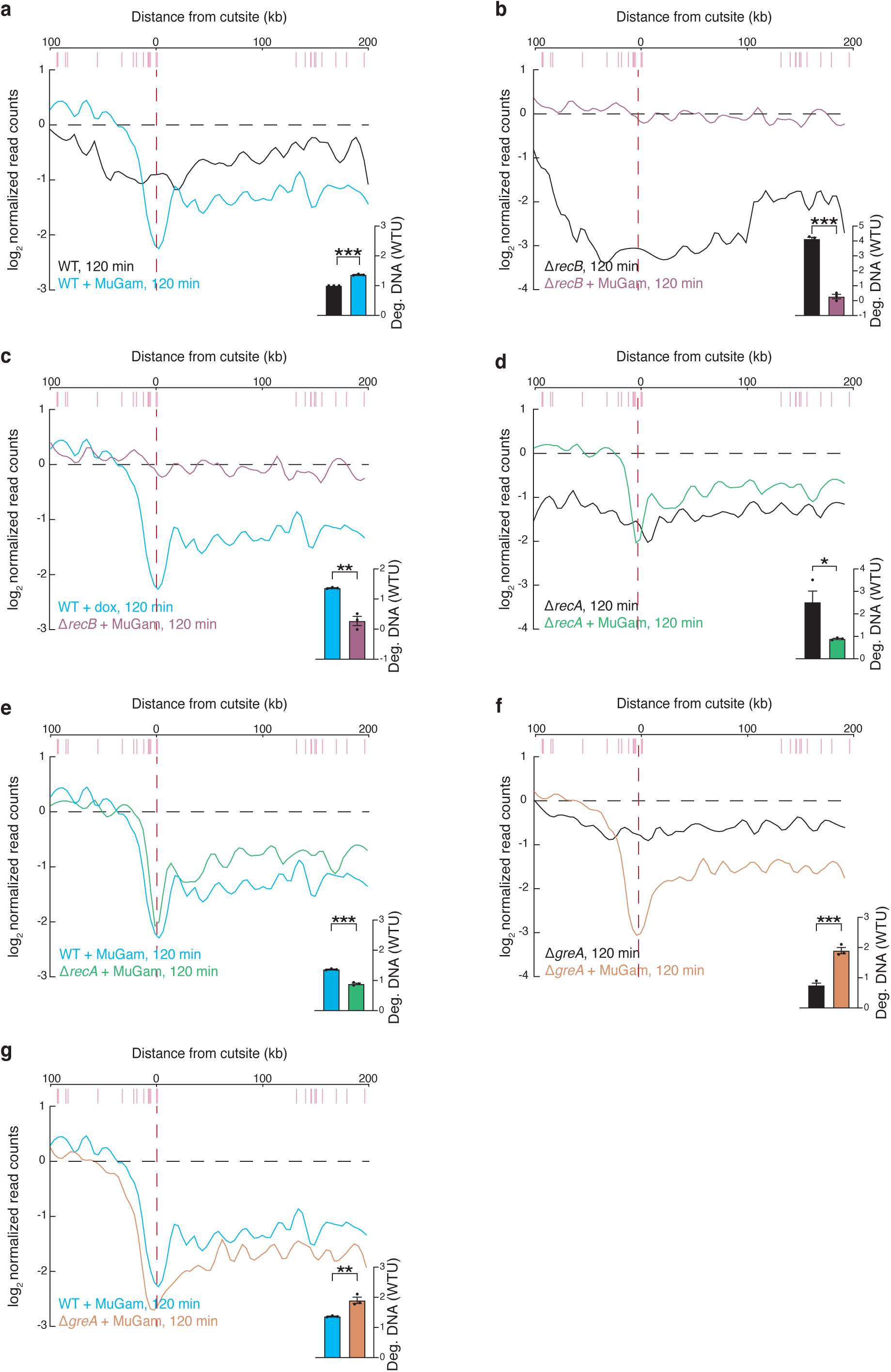
DNA degradation around the DBS site is altered by MuGam binding. **a-f** XO-seq degradation in the indicated mutants with or without MuGam expression (100 ng/ml dox) 120 minutes after I-SceI induced DSB induction at *lacA*. Red vertical line indicates position of DSB site. Pink lines show Chi site positions. Representative graph shown, bar plots are DNA degradation as measured by area under the curve, mean ±S.E.M. \*\*\**P* ≤ 0.001, **P≤ 0.01, ≤ 0.05 (two-tailed two sample *t*-test).

To study the role of MuGam on DNA resection during DSB repair, we performed XO-seq after inducing DSBs at *lacA* in cells expressing MuGam. Under these conditions, we observed greater depletion of reads (DNA degradation) around the cutsite when compared to WT cells not expressing MuGam (Fig. 3a). A deeper 20 Kb canyon is formed around the break site in presence of MuGam compared to no MuGam, likely due to the inability of cells to repair the DNA break and reconstitute the chromosome (Fig. 3a).

To determine if RecBCD is responsible for the resection around the break, we performed a similar experiment in a Δ*recB* mutant. In the absence of MuGam, degradation in Δ*recB* cells is more extensive compared to WT cells (Fig. 3a, 3b). Other exonucleases such as RecJ likely drive the observed resection^19^. When MuGam is expressed in Δ*recB* cells, strikingly, no DNA degradation around the cutsite was observed upon break induction (Fig. 3b, 3c), suggesting that MuGam-bound ends are protected from other bacterial exonucleases. These data lend further support to a model where RecBCD can process DNA ends in the presence of MuGam, albeit at a much-reduced efficiency^4^. Importantly, while Δ*recB* bacteria showed much reduced resection in the presence of MuGam, this did not translate to enhanced survival over WT cells (Fig. 2f).

To explore how the slow-down of resection by MuGam in turn affects RecA loading, we next performed XO-seq in Δ*recA* cells. Without MuGam, Δ*recA* cells display extensive DNA degradation, extending over 1 Mb downstream of the I-SceI site (Fig. 3d)^19^. In the presence of MuGam, DNA resection was significantly impeded in Δ*recA* mutants, and was even lower than in WT cells expressing MuGam (Fig. 3e). This supports the model proposed by Battacharya et al., in which MuGam can slow down RecBCD resection. Even in the absence of RecA-mediated recombination, further degradation of DNA is still prevented by MuGam.

To investigate the mechanism by which GreA impedes DSB repair at MuGam-bound DNA ends, we performed XO-seq in the Δ*greA* mutant and observed a significant increase in degradation compared with WT cells (Fig. 3g). This suggests GreA slows resection only when MuGam is present. In the absence of MuGam, however, Δ*greA* cells show reduced resection as has been previously reported (Fig. 3f)^22^.

### MuGam is retained at the DNA end in the presence GreB

Based on the above results, we hypothesize that when an RNAP complex encounters MuGam in the absence of GreA, it may displace the end-binding protein from the DNA. To investigate RNAP-MuGAM interactions, we performed *in vitro* transcription experiments with purified MuGam. As both Gre factors have similar effects on RNA synthesis and similar MuGam-related survival phenotypes, *in vitro* assays were performed with GreB, which binds RNAP with higher affinity^35^. Transcription was initiated on a linear end-labelled DNA template using biotin-tagged RNAP immobilized on streptavidin-coated magnetic beads. Synchronized halted transcription elongation complexes were formed with an incomplete set of NTP substrates. MuGam was then added, and transcription was restarted upon addition of all NTPs. RNAP dissociation from DNA was followed by measuring the DNA release into the supernatant (Fig. 4a). In the absence of MuGam, RNAP is expected to run-off the DNA end, and 50% of DNA was released after 12 min, (Fig. 4b); the remaining DNA is likely sequestered in nonproductive complexes. By contrast, DNA release was slowed in the presence of MuGam (Fig. 4b), suggesting that MuGam traps RNAP at the DNA end. Transcript cleavage factors increase torsional capacity of RNAP^36^ and can promote transcription through roadblocks, such as nucleosome^37^. Strikingly, however, while DNA dissociation was unaffected by GreB alone (Fig. 4c), when both MuGam and GreB were present, more RNAP was retained (Fig. 4d). The RNA cleavage-stimulation activity of GreB requires two acidic amino acids at the tip of its extended coiled-coil domain^29^. Substitution of one of these residues, aspartic acid 41, to asparagine impairs GreB activity^29^. The GreB D41N protein is thus unable to revive backtracked complexes. *In vitro*, DNA release over time was comparable with and without GreB D41N addition (Fig. 4d). Since RNAP retention stimulated by GreB was only observed when MuGam was bound to DNA, and the effect was abrogated when the anti-backtracking activity of GreB was impaired, we conclude that RNAP backtracks when it encounters MuGam bound to DNA (Fig. 4e). Without Gre factors present to reverse backtracking, RNAP may be able to evict MuGam, allowing for DSB repair to ensue. In the presence of Gre factors, a futile cycle of transcription restart and hitting a MuGam roadblock could potentially inhibit DSB repair (Fig. 4e).

**Figure 4.**
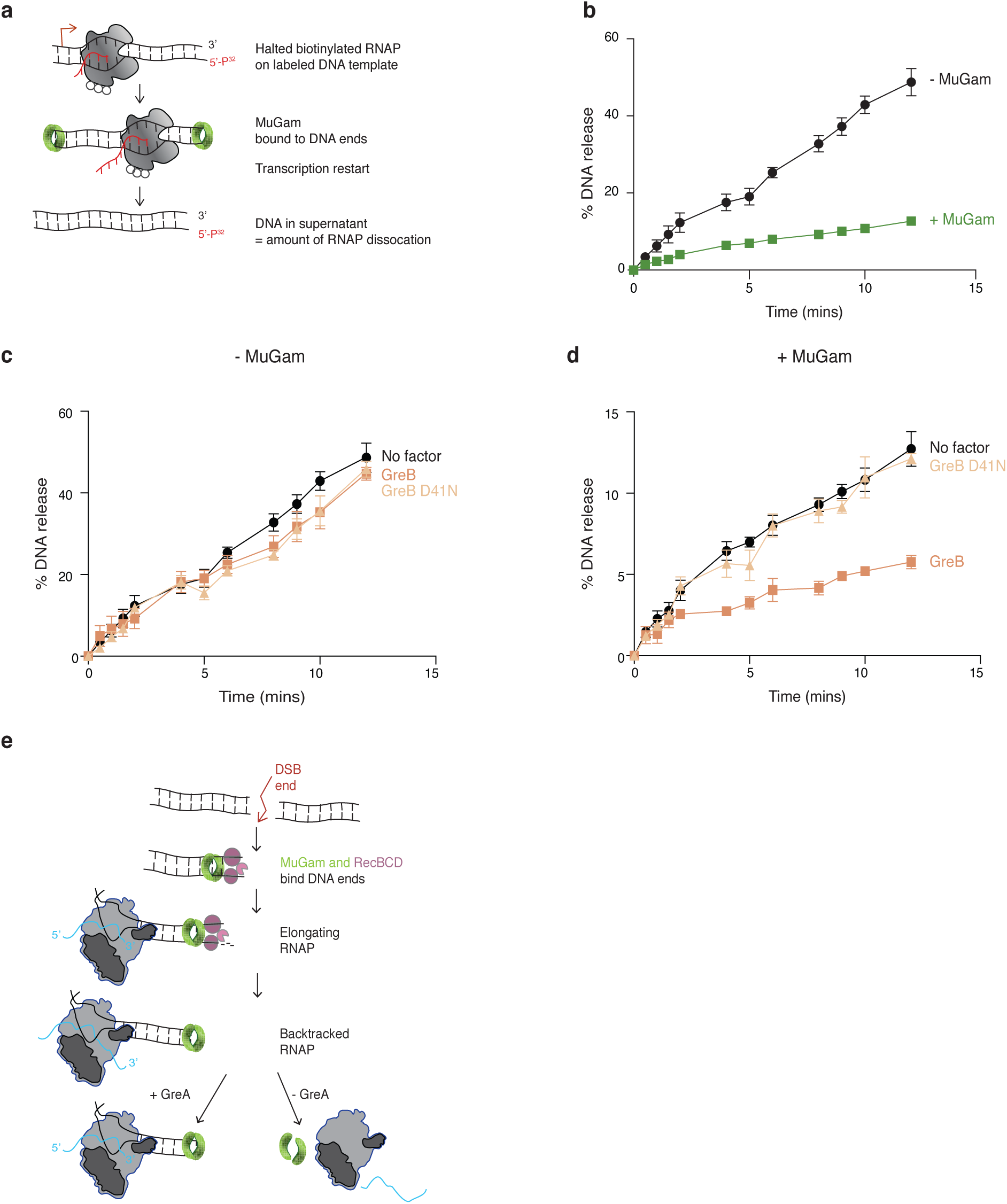
MuGam remains bound at the DNA end in the presence of the RNAP secondary channel interactor GreB. **a** Schematic of the *in vitro* experiment (see methods). **b-d** *In vitro* measurements of DNA release from the immobilized RNAP over time in the presence or absence of indicated factors. **e** Model of RNAP, MuGam and DSB repair interaction. MuGam binds to DSB ends and slows down RecBCD resection. RNAP backtracks when it encounters a protein obstacle on the DNA (MuGam). In the presence of GreA, a futile cycle of backtracking and restart ensues, preventing DSB repair. Without GreA, RNAP and MuGam are removed from the DNA, allowing for normal DSB repair.

### Subinhibitory concentration of doxycycline induces DNA breaks

While testing different dox concentrations to induce MuGam, we observed that 200 ng/ml of dox dramatically reduced viability in WT cells compared to a lower dose of 100 ng/ml, but only in strains containing the dox-inducible MuGam (Fig. 5a). WT cells lacking MuGam were unaffected at either the 100 or 200 subinhibitory antibiotic dose, as expected (Fig. 5a). We also observed that this dose-dependent toxicity is almost completely suppressed in the *greA* deletion mutant (Fig. 5a). Western blot analysis revealed that levels of MuGam protein are not increased at higher concentrations of dox (100 ng/ml vs. 200 ng/ml and 400 ng/ml) (Supplemental Fig. 1a), suggesting that dox and MuGam elicit a synergistic toxicity at these doses. We hypothesized that at higher doses, dox was contributing to the phenotype by inducing the formation of DNA breaks. To examine DNA break formation, we visualized the formation of MuGam-GFP foci at different concentrations of dox. We found that the number of WT cells with MuGam-GFP foci increases with dox dosage, and ∼40% of cells have at least one MuGam-GFP focus at the highest dose of 400 ng/ml dox (Fig. 5b). This suggests that even at subinhibitory doses of dox, more DNA breaks are induced in WT cells.

**Figure 5.**
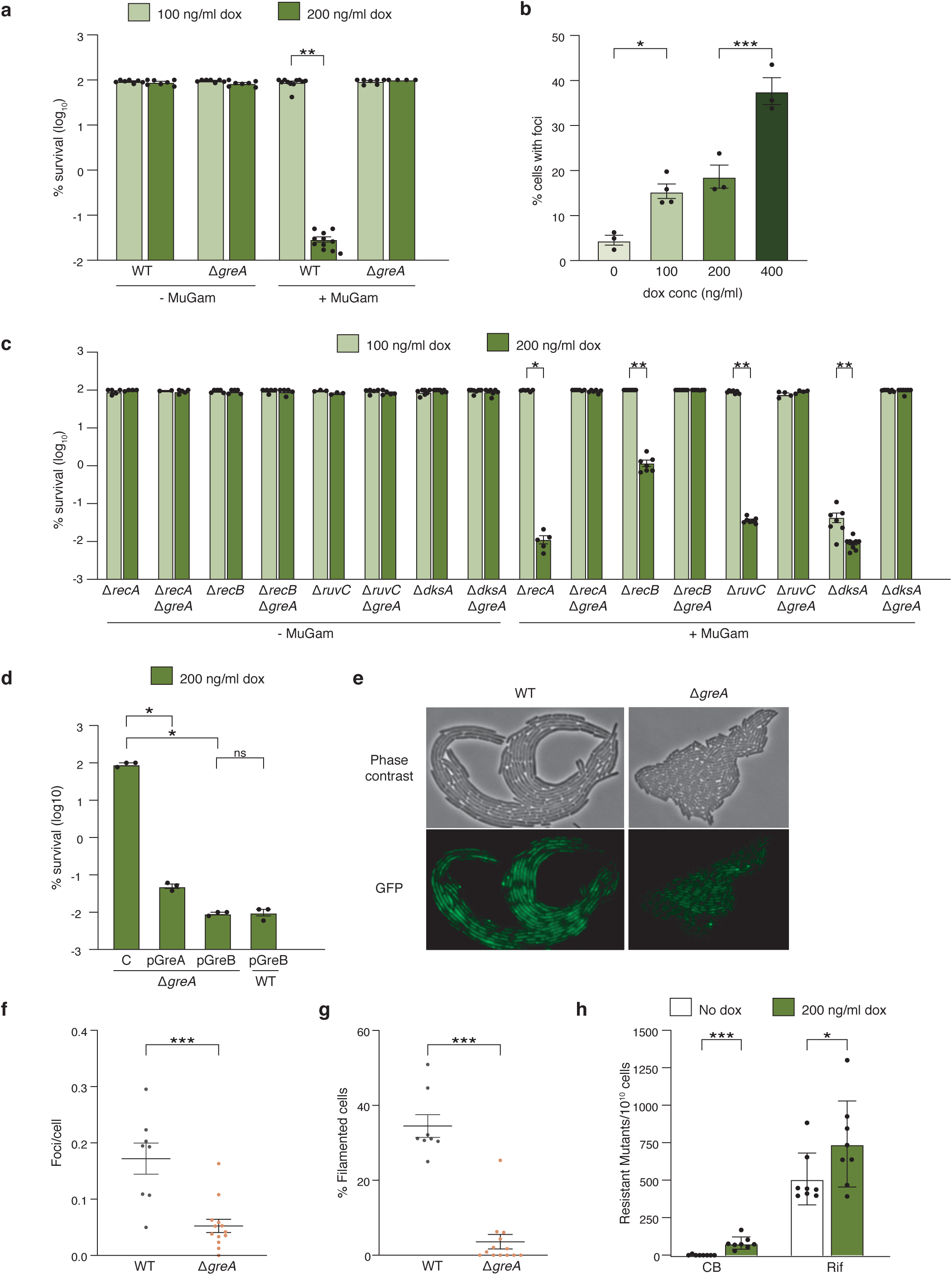
High dose of dox induces DSBs, which is reduced in the absence of GreA. **a** Quantitative survival of WT and the Δ*greA* mutant strain at low (100 ng/ml) and high (200 ng/ml) dox; mean ±S.E.M. \*\**P* ≤ 0.01 (Kruskal–Wallis test followed by pairwise Wilcox test with multiple testing correction). **b** Quantification of MuGam-GFP foci in WT cells grown in the presence of increasing concentrations of dox; mean ±S.E.M. \*\*\**P* ≤ 0.001, \**P* ≤ 0.05 (One-way ANNOVA, Tukey Posthoc). **c** Quantitative survival of the indicated mutants at either 100 ng/ml or 200 ng/ml dox; mean ±S.E.M. \*\**P* ≤ 0.01, \**P* ≤ 0.05 (Kruskal–Wallis test followed by pairwise Wilcox test with multiple testing correction). **d** Quantitative survival after expression of control plasmid (c) or high-copy plasmids expressing either GreA or GreB (0.1 mM IPTG induction) in the indicated mutants, mean ±S.E.M. \**P* ≤ 0.05. **e** Bright-field and GFP images of actively growing WT and Δ*greA* microcolonies on agar pads containing 400 ng/ml dox followed by time-lapse microscopy. **g,f** Percentage of filamented cells (> 6 μM) and quantification of MuGam-GFP foci from e; n ≥ 3 microcolonies (dots represent individual microcolonies analyzed), mean ±S.E.M. \*\*\*\**P* < 0.0001, \*\*\**P* = 0.0002 (unpaired t-test). **h** Carbenicillin (CB) and rifampicin (Rif) mutation rate, n ≥ 3 biological replicates, mean ±S.E.M. \*\*\**P* ≤ 0.001, \**P* ≤ 0.05 (Welch’s one-sided t-test).

Unlike I-SceI- or PHL-induced DSBs, the Δ*greA*-mediated suppression of dox-induced DSB repair was not RecA or RecB dependent, indicating mechanisms and proteins that are not involved in the canonical recombination DSB break repair pathway may play a role in this phenotype (Fig. 5c). Since spontaneous DSBs are thought to largely arise during replication stalling and have been shown to require the RuvC helicase for processing of recombination intermediates^38^, we tested the Δ*ruvC* mutant for dox-induced toxicity. We found that this mutant was as sensitive to dox as the WT strain, arguing against RuvC involvement in the Δ*greA*-mediated mediated suppression (Fig. 5c).

To study whether both Gre factors act similarly in the mechanism involved in dox-induced breaks, GreA and GreB were overexpressed from plasmids in both WT and *ΔgreA* backgrounds. Both Gre factors when overexpressed abrogate the resistance of the Δ*greA* mutant in the presence of MuGam, suggesting that breaks induced by dox are affected by RNAP backtracking, similar to PHL-induced breaks (Fig. 5d). Interestingly, compared with WT, the Δ*dksA* strain was ∼ 2000-fold more sensitive to MuGam even at the lowest dose of dox (100 ng/ml) (Fig. 5c). However, it was rescued by the deletion of *greA* (Fig. 5c), consistent with a model in which DksA promotes cell survival by excluding GreA from RNAP (Fig. 5d). Indeed, overexpression of super DksA mutants that have higher affinity for RNAP^39^ rescues WT sensitivity to 200 ng/ml dox (Supplementary Fig. 1b).

The above experiments may implicate a transcription-mediated mechanism in the formation of dox-induced DSBs that is independent of the canonical HR pathway. We have previously shown that uncoupling transcription and translation in *E. coli* results in increased transcription-replication conflicts^23^. Since dox is known to affect active ribosomes it is possible that DSBs at higher dox concentrations are a result of an uncoupling between translation and transcription^40^. To look more closely at the formation of DSBs during active growth *in vivo*, we grew MuGam-GFP expressing WT and Δ*greA* microcolonies on high dose (400 ng/ml) dox-containing agar pads and performed time-lapse fluorescence microscopy to determine the appearance and level of MuGam-GFP foci in individual cells (Fig. 5e). We found that WT cells contained significantly more MuGam-GFP foci than Δ*greA* cells, indicating that fewer DNA breaks form in the absence of GreA. ∼30% of WT cells formed elongated filaments, whereas Δ*greA* cells produced very few filaments (Fig. 5g). Cell division arrest leading to filamentation is a classic phenotype indicative of DSB-dependent DNA damage (SOS) response^41^. Reduced filamentation in Δ*greA* cells suggests that the SOS response in not induced, potentially because fewer DSB breaks occur at high dox concentrations in these mutants (Fig. 5f). Given that transcriptional arrest tends to occur in the absence of GreA, and that a high dose of dox likely slows down translation, the respective rates of these processes may be better matched in Δ*greA* mutants, improving transcription-translation coupling, thereby reducing the formation of DSBs compared to WT cells under these conditions.

A potential consequence of increased DSBs is an increase in mutation rate^42^. At the high dox concentrations, we found that resistance to the antibiotics carbenicillin and rifampicin was increased, implicating dox-induced DSBs in mutagenesis (Fig. 5h).

## DISCUSSION

Proteins involved in DNA replication, repair, and transcription share the same DNA template and coordination between these processes is indispensable in preventing conflicts, which could result in genomic instability. Studies in *E. coli* have pinpointed a role of RNAP secondary channel interactors in modulating transcription conflicts with replication as well as DNA repair machineries^22,43,44^. Here, we investigate how the modulation of RNAP processivity influences DSB repair in the presence of a DNA end-binding protein, MuGam.

Our study confirms that the expression of MuGam results in reduced bacterial viability as previously observed in WT cells with and without I-SceI-induced genomic damage; and extends this effect to DSBs induced by phleomycin^8^. This reduced survival was initially thought to be a consequence of MuGam blocking RecBCD access to double-strand DNA ends (DSEs). Recent *in vitro* data indicates, however, that MuGam does not completely block RecBCD activity but rather reduces its processivity^4^. MuGam dimer, shaped as a donut, is reported to bind DNA ends and slide along DNA *in vitro*, propelled by RecBCD translocase activity^4^. Here, using XO-seq to directly examine resection at I-SceI-induced DSBs in live cells, we found that RecB-mediated DNA resection was indeed influenced by the presence of MuGam. Furthermore, in the absence of RecB, MuGam binding at DNA-ends appears to effectively block all other exonucleases from acting on DSEs (Fig. 3b). In striking contrast to the extensive DNA degradation after I-SceI cutting seen in Δ*recA* mutants not expressing MuGam, when MuGam is present, Δ*recA* cells show less degradation around the I-SceI site (Fig. 3d). This suggests that MuGam alone may be sufficient to reduce RecBCD processivity. Altogether, our data shows that MuGam inhibits break repair *in vivo* by altering RecBCD resection kinetics and not by just shielding DNA ends from RecBCD as previously thought. We find that DNA resection by RecBCD in the presence of MuGam can go as far as ∼30 kb, passing multiple Chi sites without promoting break repair. This suggests that in addition to its effects on RecBCD processivity, MuGam may also inhibit other functions of RecBCD important for recombination, such as exonuclease polarity switching activity or RecA loading.

One key finding in our study is that RNAP processivity influences DSB repair when MuGam is expressed in *E. coli*. Factors that rescue RNAP backtracking, such as GreA and GreB, increase sensitivity to MuGam (Fig. 2). Counterintuitively, the increased resistance of the Δ*greA* strain to MuGam expression after DSB induction is accompanied by an increase in DNA degradation on both sides of the DSB (Fig. 3f, 3g). Our *in vitro* transcription experiments show that following MuGam binding to DSEs, RNAP release is inhibited, and this effect is exacerbated by GreB (Fig. 4c, 4d). We propose a model in which loss of Gre factors enhances dissociation of MuGam from DSBs along with RNAP, freeing the ends for efficient repair (Fig. 4e). We further speculate that ejection of MuGam from the DNA restores RecBCD activity, promoting DNA resection and RecA loading (Fig. 4e). MuGam bound to DNA ends likely serves as a substantial roadblock to transcription and may be released from the DNA by RNAP, but how MuGam is released from DNA is unclear. It remains to be determined if dissociation is passive or requires additional factors for active removal. DNA curtain assays show that MuGam dissociates only after RecBCD stops translocating, suggesting that MuGam may be trapped by the advancing exonuclease^4^. In our model, we propose that when GreA is present, futile attempts to restart transcription adjacent to the DSB impedes MuGam removal and inhibits repair (Fig. 4e).

Dox is an antibiotic used to treat a wide variety of Gram-positive and Gram-negative bacteria that cause pneumonia, sexually transmitted infections, and vector-borne diseases. We observed that subinhibitory doses of dox is toxic to WT cells expressing MuGam. Western blot analysis revealed that MuGam protein levels were not changed at higher doses of dox (Supplemental Fig. 1a), and yet DSBs occurred in a dose-responsive manner in WT cells, as measured by MuGam-GFP foci (Fig. 5b). These results lead us to conclude that a subinhibitory dose of dox is capable of inducing DSBs, which increases resistance to other antibiotics (Fig. 5h), possibly via the stress-induced mutagenesis pathway^45^.

The effects of dox-induced DSBs are only revealed when MuGam is expressed, leading to a synergistic loss of survival, which is surprisingly not dependent on the canonical RecA-mediated HR pathway (Fig. 5c). We show that the synergistic loss of survival is completely rescued by loss of GreA. The presence of Gre factors and co-translating ribosomes have been shown to suppress RNAP backtracking^46^. Since higher doses of dox are known to affect translation, and translation and transcription are coupled in *E. coli*, the rescue of this toxicity in Δ*greA* cells implicates transcription and/or backtracked RNAP in this mechanism of DSB generation. Importantly, we found that Δ*greA* cells have fewer DSBs compared to WT (Fig. 5e, 5f) at higher dox doses as measured by MuGam-GFP foci. A likely DNA-end target for MuGam under these conditions could be stalled replication forks^8^. Fewer stalled replication forks could be occurring in Δ*greA* cells due to the presence of DksA, which was previously shown to ameliorate transcription:translation uncoupling during amino acid starvation by releasing RNAP^21,23^. Therefore, in Δ*greA dksA+* cells, the presence of DksA may release most backtracked RNAP and allow trailing ribosomes to catch up with transcription leading to better translation-transcription coupling. This new finding raises important questions about how an antibiotic targeting the ribosome may trigger DNA instability involving replication/transcription conflicts. Additional studies will be required to fully address this important issue and dissect the mechanism involved.

In a broader context, this work raises questions about the interaction between Ku, the MuGam homolog, and transcription at DSBs in eukaryotes. We have observed that GreA impedes DSB repair regardless of whether the DNA end is free or bound by MuGam. Perhaps similar interactions occur between TFIIS and Ku in higher organisms, reiterating the importance of understanding how transcription processivity influences DSB repair and the mechanisms that reduce conflicts and maintain genomic stability in growing cells.

## Acknowledgements

We thank Susan Rosenberg for sharing strains and reagents. The study was supported by the NIH grant DP1-AI152073 (C.H) and R01-R01GM067153 (I.A).

---

**Supplementary Figure 1.**
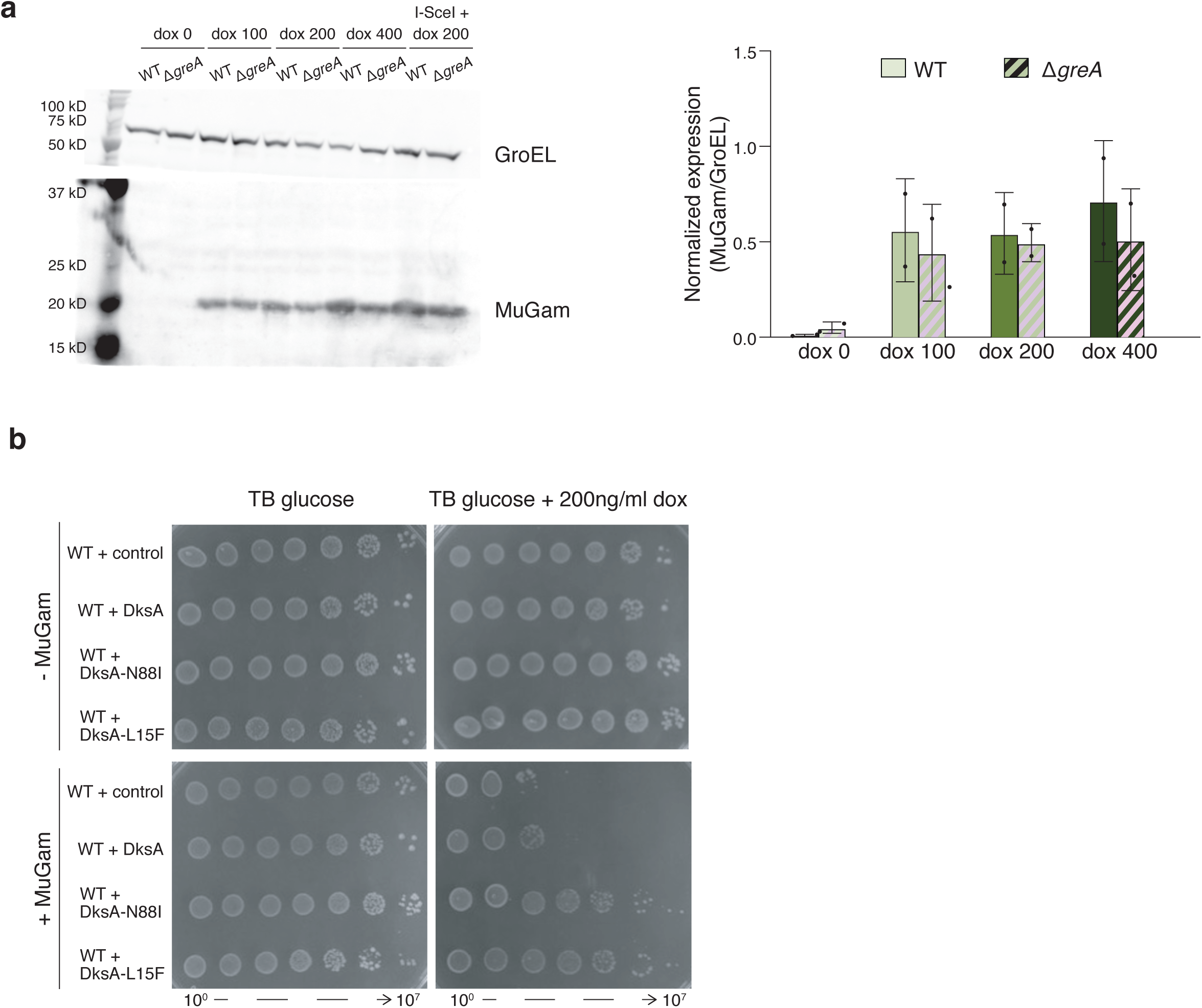
Higher doses of dox does not increase MuGam levels and overexpression of super DksAs rescues survival at high dox concentrations. **a** Quantification of Western blot for MuGam after incubation of WT and Δ*greA* cells with 0, 100, 200, and 400 ng/ml dox using GroEL for normalization of and quantification of normalized protein levels, mean ±S.E.M. **b** Semiquantitative survival assays in strains with or without MuGam (dox 100ng/ml) containing control plasmid or plasmids overexpressing super DksA mutants (N88I and L15F) (0.1 mM IPTG).

