## Supplementary tables for "Modulation of RNA polymerase processivity affects double-strand break repair in the presence of a DNA end-binding protein"

**Supplementary Table 1. Bacterial strains**

| Strain number | Genotype | Reference |
|---|---|---|
| CH30 | Wild-type MG1655 | Lab stock |
| CH393 | Δ*greA*::FRT | Lab stock |
| CH413 | Δ*greA*::FRTKanFRT | Lab stock |
| CH580 | Δ*dksA*::FRTKanFRT | Lab stock |
| CH618 | Δ*dksA*::FRTKanFRT<br>Δ*greA*::FRT | Lab stock |
| CH1039 | Wild-type MG1655<br>+[pBA169Amp] | CH30+ pBA169Amp |
| JW2788 | Δ*recB*::FRTKanFRT | JW2788<br>Baba et al., 2006 |
| CH3782 | Δ*araBAD567* Δ*attλ*::$P_{BAD}$<br>*zfd2509.2*::$P_{N25}$*tetR*FRT<br>Δ*attTn7*::FRT*cat*FRT $P_{N25}$-*tetO-*<br>*gam-gfp* | SMR 14334<br>Shee et al., 2003 |
| CH3788 | Δ*araBAD567* Δ*attλ*::$P_{BAD}$<br>*zfd2509.2*::$P_{N25}$*tetR*FRT<br>Δ*attTn7*::FRT*cat*FRT $P_{N25}$-*tetO-*<br>*gam-gfp* Δ*greA*::FRTKanFRT | CH3782 X P1(CH413) |
| CH4088 | Δ*recA*::FRTKanFRT | Lab stock |
| CH4199 | Δ*attλ*::$P_{N25}$*tetR*FRT<br>Δ*attTn7*::FRT*cat*FRT<br>$P_{N25}$-*tetO-gam* Strep | SMR13974, from S.M.<br>Rosenberg |
| CH4200 | FC40 *yfeP-*<br>$P_{N25}$*tetR*::FRTKanFRT | Shee et al., 2013 |
| CH4230 | *yfeP*-$P_{N25}$*tetR*::FRTKanFRT | CH30 X P1(CH4200) |
| CH4242 | *yfeP*-$P_{N25}$*tetR*::FRTKanFRT<br>Δ*attTn7*::FRT*cat*FRT<br>$P_{N25}$-*tetO-gam* | CH4230 X P1(CH4199) |
| CH4252 | *yfeP*-$P_{N25}$*tetR*::FRT<br>Δ*attTn7*::FRT $P_{N25}$-*tetO-gam* | CH4242 flipped |
| CH4272 | *yfeP*-$P_{N25}$*tetR*::FRT<br>Δ*attTn7*::FRT $P_{N25}$-*tetO-gam*<br>Δ*greA*::FRTKanFRT | CH4252 X P1(CH413) |
| CH4782 | Δ*ruvC*::FRTKanFRT | Lab stock |
| CH4784 | Δ*greA*::FRT *ruvC*::FRTKanFRT | CH393 X P1(CH4782) |
| CH4945 | *yfeP*-$P_{N25}$*tetR*::FRT<br>$P_{N25}$-*tetO*-gam Δ*greA*::FRT | CH4272 flipped |
| CH5069 | *yfeP*-$P_{N25}$ *tetR*::FRT<br>$P_{N25}$-*tetO*-gam | CH4252 X P1(CH580) |

| | | |
|---|---|---|
| | Δ*dksA*::FRTKanFRT | |
| CH5079 | *yfeP*-$P_{N25}$ *tetR*::FRT $P_{N25}$-*tetO*-gam Δ*greA*::FRT Δ*dksA*::FRTKanFRT | CH 4945 X P1(CH580) |
| CH5097 | Δ*araBAD567* Δattλ::$P_{BAD}$ Zfd2509.2::$P_{N25}$*tetR*::FRT ΔattTn7::FRT*cat*FRT $P_{N25}$-tet-O-gam GFP *recB*::FRTKanFRT[pNT3*recB*] | CH 3929 +pNT3recB Amp |
| CH5100 | Δ*recB*::FRTKanFRT | CH30 X P1(CH5097) |
| CH5102 | Δ*greA*::FRT Δ*recB*::FRTKanFRT | CH393 X P1(5097) |
| CH5120 | *yfeP*-$P_{N25}$ *tetR*::FRT $P_{N25}$-*tetO*-gam Δ*recA*::FRTKanFRT | CH4252 X P1(CH4088) |
| CH5122 | *yfeP*-$P_{N25}$ *tetR*::FRT $P_{N25}$-*tetO*-gam Δ*greA*::FRT Δ*recA*::FRTKanFRT | CH4945 X P1(CH4088) |
| CH5123 | *yfeP*-$P_{N25}$ *tetR*::FRT $P_{N25}$-*tetO*-gam Δ*recB*::FRT*kan*FRT | CH4252 X P1(5097) |
| CH5125 | *yfeP*-$P_{N25}$ *tetR*::FRT $P_{N25}$-*tetO*-gam Δ*recB*::FRTKanFRT Δ*greA*::FRT | CH4945 X P1(5097) |
| CH5236 | Δ*recA*::FRT | Lab stock |
| CH5238 | Δ*greA*::FRT Δ*recA*::FRT | Lab stock |
| CH6791 | Wild-type MG1655 +[pBA169DksA Amp] | CH30+pBA169DksA Amp |
| CH7058 | Δ*attλ*::$P_{BAD}$-ISce1 Δ*attTn7*::FRT $P_{N25}$-*tetO-gam yfeP*-$P_{N25}$*tetR* FRT *lacA*::I-SceI FRT | Lab stock |
| CH7063 | Δ*attλ*::$P_{BAD}$-ISce1 Δ*attTn7*::FRT $P_{N25}$-*tetO-gam yfeP*-$P_{N25}$*tetR* FRT *lacA*::I-SceI FRT Δ*greA*::FRT | CH7058 X P1(413) flipped out |
| CH7093 | Δ*attλ*::$P_{BAD}$-ISce1 Δ*attTn7*::FRT $P_{N25}$-*tetO-gam yfeP*-$P_{N25}$*TetR* FRT *lacA*::I-SceI FRT Δ*recB*::FRTKan FRT | CH7058 X P1(5097) |
| CH7094 | Δ*attλ*::$P_{BAD}$-ISce1 Δ*attTn7*::FRT $P_{N25}$-*tetO-gam yfeP*-$P_{N25}$*tetR* FRT *lacA*::I-SceI FRT Δ*recB*::FRTKanFRTΔ*greA*::FRT | CH7063 X P1(5097) |

| | | |
|---|---|---|
| CH7100 | Δ*attλ*::$P_{BAD}$-ISce1 Δ*attTn7*::FRT $P_{N25}$-*tetO-gam yfeP*-$P_{N25}$*tetR* FRT *lacA*::I-SceI FRT Δ*recA*::FRTKanFRT | CH7058 X P1(4088) |
| CH7102 | Δ*attλ*::$P_{BAD}$-ISce1 Δ*attTn7*::FRT $P_{N25}$-*tetO-gam yfeP*-$P_{N25}$*tetR* FRT *lacA*::I-SceI FRT Δ*recA*::FRTKanFRT Δ*greA*::FRT | CH7063 X P1(4088) |
| CH7104 | Δ*attλ*::$P_{BAD}$-ISce1 Δ*attTn7*::FRT $P_{N25}$-*tetO-gam yfeP*-$P_{N25}$*tetR* FRT *lacA*::I-SceI FRT Δ*greA*::FRTΔ*dksa*::FRTKanFRT | CH7063 X P1(580) |
| CH7106 | Δ*attλ*::$P_{BAD}$-ISce1 Δ*attTn7*::FRT $P_{N25}$-*tetO-gam yfeP*-$P_{N25}$*tetR* FRT *lacA*::I-SceI FRT Δ*dksa*::FRTKanFRT | CH7058 X P1(580) |
| CH7437 | Δ*attλ*::$P_{BAD}$-ISce1 Δ*attTn7*::FRT $P_{N25}$-*tetO-gam yfeP*-$P_{N25}$*tetR* FRT *lacA*::I-SceI FRT [pBA169 Amp] | CH7058 +pBA169 Amp |
| CH7439 | Δ*attλ*::$P_{BAD}$-ISce1 Δ*attTn7*::FRT $P_{N25}$-*tetO-gam yfeP*-$P_{N25}$*tetR* FRT *lacA*::I-SceIFRT [pBA169*greB* Amp] | CH7058+pBA169*greB* Amp |
| CH7440 | Δ*attλ*::$P_{BAD}$-ISce1 Δ*attTn7*::FRT PN25-*tetO-gam yfeP*-$P_{N25}$*tetR* FRT *lacA*::I-SceIFRT Δ*greA*::FRT[pBA169 Amp] | CH7063+pBA169 Amp |
| CH7442 | Δ*attλ*::$P_{BAD}$-ISce1 Δ*attTn7*::FRT PN25-*tetO-gam yfeP*-$P_{N25}$*tetR* FRT *lacA*::I-SceI FRT Δ*greA*::FRT[pBA169*greB* Amp] | CH7063+ pBA169*greB* Amp |
| CH7607 | Δ*ruvABC*::*cat* [pGB*ruvABC* Spec] | SMR8899 from S.M. Rosenberg lab |
| CH7653 | *yfeP*-$P_{N25}$*tetR*::FRT Δ*attTn7*::FRT $P_{N25}$-*tetO--gam* Δ*ruvABC*::*cat* | CH4252 X P1(7607) |
| CH7655 | *yfeP*-$P_{N25}$-*tetR*::FRT Δ*attTn7*::FRT $P_{N25}$-*tetO--gam* Δ*ruvC*::*cat* Δ*greA*::FRTKanFRT | CH4272 X P1(7607) |
| CH9411 | CH4252+[pBA169 Amp] | CH4252+ pBA169 Amp |

| | | |
|---|---|---|
| CH9417 | Wild-type MG1655 + [pBA169DksA-N88I Amp] | CH30+ pBA169DksA-N88I Amp |
| CH9419 | Wild-type MG1655 +[pBA169DksA-L15F Amp] | CH30+ pBA169DksA-L15F Amp |
| CH9421 | CH4252+[pBA169DksA Amp] | CH4252+ pBA169DksA Amp |
| CH9423 | CH4252+[pBA169DksA-L15F Amp] | CH30+ +[pBA169DksA-L15F] |

**Supplementary Table 2. Plasmids**

| Plasmid | Genotype | Reference |
|---|---|---|
| pBR322 | Common vector backbone<br>Amp | Lab stock |
| pBR322-GreB | *greB* | Satory et al., 2013 |
| pET-28a<br>N-ter His-tag Mu Gam | *lacUV5*::*t7 rnap*<br>$P_{T7}$ *gam*<br>*6Xhis cat* Kan | Rosenberg lab gift |
| p-NT3 | $P_{tac}$ *recB* Amp | Saka et al., 2005<br>Lab stock |
| pBA169 | Common vector backbone | Lab stock |
| pBA169-GreA | *greA* | Lab stock |
| pBA169-GreB | *greB* | Lab stock |
